# Shared Pain, Shared Decisions: How Empathy Shapes Social Conformity Through Physiology and Visual Attention

**DOI:** 10.64898/2026.06.27.734940

**Authors:** Soroosh Golbabaei, Martin Walter, Khatereh Borhani

## Abstract

Empathy enables individuals to attune to others’ experiences through shared affective, sensorimotor, and neural representations, but its influence on higher-level decision-making and social alignment remains unknown. We examined whether empathy promotes social conformity and whether this effect depends on physiological arousal, visual attention, and socio-emotional traits. Seventy-three participants completed an empathy-for-pain task followed by a modified random-dot task in which they interacted with previously seen empathy targets, while physiological signals and eye movements were recorded. Drift-diffusion modeling showed that feedback from empathy targets increased conformity, reflected in stronger decision bias toward targets, faster evidence accumulation, and reduced decision conservatism. Visual attention to the eyes or mouth was linked to greater conformity depending on the context. Heart rate variability was unrelated to conformity, whereas phasic skin conductance was associated with smaller bias changes and greater conservatism. Alexithymia and autistic traits further shaped the empathy-conformity relationships. These findings suggest that empathic shared representation extends beyond affective resonance to influence social alignment in decision-making.

---

Empathy is a multifaceted process through which individuals share, understand, and care about the internal states of others ^1,2^. It involves several partially dissociable components, including affective resonance with another person’s sensorimotor and emotional states, often described as emotional contagion or affective empathy; cognitive understanding of another person’s feelings, thoughts, motivations, and intentions; and prosocial motivation to respond to another person’s needs, such as helping to alleviate their pain ^3–5^. Importantly, empathic engagement is not merely a matter of physiological or neural alignment. Rather, empathy may also alter how the self is represented in relation to another person ^6–8^. A growing body of work suggests that empathic engagement can reduce the perceived boundary between self and other, producing what is commonly referred to as self–other overlap or reduced self–other distinction ^7,9,10^.

This idea is also compatible with predictive coding accounts of social cognition. According to predictive coding frameworks, the brain and body continuously generate probabilistic predictions about the self, bodily states, sensory input, emotional experience, and the surrounding social environment ^11–13^. Incoming information is evaluated against these predictions, and discrepancies between predicted and observed states generate prediction errors that the system attempts to minimize. In the context of empathy, resonating with another person’s emotional or sensorimotor state may introduce a discrepancy between the predicted representation of the self as distinct from others and the incoming representation of another person’s state ^14,15^. Reducing this discrepancy may require partial alignment between self and other, thereby increasing self–other overlap. In this sense, empathy may be understood as a process through which the self becomes temporarily attuned to another person’s internal state.

A recent theoretical account has suggested that such empathic alignment may extend beyond affective and motor synchrony to broader cognitive and social alignment, with greater alignment emerging among ingroup members and individuals who feel closer to one another^16^. Although this possibility has not yet been directly examined, indirect evidence is consistent with it. For example, empathic engagement can reduce perceived interpersonal physical distance ^7^, increase identification with others ^17,18^, reduce bias toward outgroups, and attenuate racial bias ^8,19^. Moreover, empathy has been associated with physiological and neural synchrony, both at the state level and as a function of trait empathy ^20–23^. Outside the empathy literature, interpersonal synchrony has been shown to promote cooperation ^24–26^, cohesion ^25,27^, friendship formation ^28^, social adjustment ^29^, imitation ^30^, perceptual conformity ^31^, and compliance with requests ^32^. Taken together, these findings raise an important but unexamined question of whether empathy also increases conformity to another person’s decisions, a question that the present study aims to address.

Conformity, or social alignment, describes the process through which individuals modify their beliefs, judgments, or behaviors in response to information from another person or group ^33–35^. Conformity can serve adaptive social and informational functions. It may facilitate social bonding ^36^, improve decisions when others possess useful information ^37,38^, reduce interpersonal conflict ^39^, and protect individuals from rejection or disapproval ^40,41^. At the same time, conformity can also be driven by normative motives, such as the desire to gain approval, avoid social exclusion, or reduce negative affect when outcomes are uncertain or unfavorable ^42^.

The link between empathy and conformity is further suggested by recent discussions of emotional conformity ^43^. In fact, emotional convergence in empathic contexts has been described as a form of emotional conformity, whereby the empathizer uses emotional information conveyed by the target, such as facial expressions or bodily signals, to align with the target’s emotional state. Another person’s emotional expression is not only something to be shared or understood. It is also social information that can shape the observer’s responses. If empathy promotes emotional alignment through the processing of another person’s affective signals, it may also promote higher-level forms of social alignment, including conformity to that person’s decisions or judgments. This possibility is consistent with accounts of conformity based on social categorization and self-categorization, according to which disagreement with those perceived as part of the self or ingroup creates pressure toward alignment ^44–46^. Because empathy reduces self–other distinction and increases perceived closeness ^47,48^, it may bring another person’s decisions into the range of information that is treated as self-relevant and therefore more likely to influence one’s own behavior.

However, conformity is not a unitary process. Recent computational approaches have shown that social influence can affect different components of decision-making ^49^. Drift diffusion models are particularly useful for separating these components. In a drift diffusion model, decisions are represented as a process of evidence accumulation over time. Information favoring each option is accumulated until it reaches a decision threshold, at which point a response is made. Social influence may affect this process in several ways. First, individuals may begin the decision process with an initial bias toward one option, reflecting prior expectations, preferences, or social information. In the context of conformity, this can be conceptualized as judgmental conformity. Second, social information may alter the rate at which evidence is accumulated, reflected in the drift rate. Depending on the task, this may represent perceptual conformity or socially biased evidence accumulation. Third, individuals may adjust how much evidence they require before making a decision, reflected in threshold separation. This parameter indexes response caution or conservativeness ^44,50^. Few previous studies have used drift diffusion models to dissociate these components of conformity ^44,50–52^. Therefore, using this model, we were interested to unpack the potential effect of empathy on different components of decision.

Empathy also depends strongly on the processing of social and emotional information, much of which is conveyed visually through facial expressions. Previous research suggests that empathy is related to how individuals allocate visual attention to others’ faces, shaped by their motivation to understand or respond to another person’s emotional state ^53^. Different emotional expressions rely on different facial regions for the communication of relevant information. In the case of pain, the eyes and mouth are especially important for conveying distress ^6,7,53,54^. In addition, visual attention has been suggested to play a role in coordination dynamics and error-correction mechanisms ^55^. Yet it remains unclear whether attention to these regions is associated with empathy-induced conformity, and if so, whether visual attention relates to judgmental bias, evidence accumulation, or decision caution.

In addition to visual attention, empathy is closely linked to physiological responding ^20,56–58^. Embodying another person’s emotional state may involve changes in bodily signals, and empathic engagement also requires the regulation of physiological arousal in response to another person’s distress ^59^. However, findings on the relationship between empathy and physiological measures such as arousal and heart rate variability have been mixed, with some studies reporting positive associations ^60–62^, others reporting null effects or negative relationships ^63^. This inconsistency highlights the need to examine whether physiological responses to others’ pain is related to the potential effect of empathy on conformity.

Individual differences may further shape the relationship between empathy and conformity. People differ in their capacity for empathic resonance, self–other overlap, physiological synchrony, and social decision-making. Two traits that are particularly relevant in this context are alexithymia and autistic traits. Both have been associated with alterations in empathy, emotional embodiment, physiological responses to emotion, and the use of feedback to update internal representations ^56,64–69^. From a predictive coding perspective, both alexithymia and autistic traits have been linked to atypical weighting of prediction errors and difficulties in integrating feedback into internal models ^66,70–72^. Alexithymia has also been associated with alterations in different components of decision-making, including initial bias, evidence accumulation, and the setting of decision boundaries ^66^. Therefore, if empathy increases conformity, alexithymia and autistic traits may moderate this effect by altering the degree to which individuals differentiate between high- and low-empathy contexts and translate empathic engagement into social alignment.

In the present study, we investigated whether empathy increases conformity to another person’s decisions and which computational components of the decision process are involved. Using a within-subject design, participants first completed an empathy-for-pain task in which high and low levels of empathy were induced toward different targets. During this task, we measured cognitive and affective empathy as well as self–other distinction. Participants then completed a cooperative random-dot motion task. On each trial, they first judged the direction of moving dots and then had the opportunity to revise their decision after receiving feedback from a confederate. Eye movements were recorded to assess visual attention to pain-relevant facial regions, particularly the eyes and mouth. Electrocardiography and skin conductance were recorded as physiological indices of arousal and autonomic regulation. Finally, participants completed the Toronto Alexithymia Scale-20 ^73–75^ and the Autism-Spectrum Quotient ^76^ to assess individual differences in alexithymia and autistic traits.

We used drift diffusion modeling to dissociate different mechanisms of conformity, including judgmental bias, evidence accumulation, and decision conservativeness. We hypothesized that participants would show stronger judgmental conformity in the high-empathy condition than in the low-empathy condition. We also explored whether empathy modulates evidence accumulation and threshold separation, reflected in drift rate and decision threshold, respectively. Although higher drift rates and lower threshold separation might be expected in line with the predicted bias effect, we had no specific hypotheses regarding these parameters. Therefore, these analyses were exploratory. In addition, we expected greater visual attention to the eyes and mouth during the empathy-for-pain task to be associated with stronger judgmental conformity, while its associations with evidence accumulation and conservativeness were examined exploratorily. Similarly, we expected physiological arousal, indexed by heart rate variability and skin conductance, to relate to judgmental conformity, with exploratory analyses examining its relationship to other decision parameters. Finally, we examined whether alexithymia and autistic traits are related to the empathy-related conformity. Based on previous work, we expected these traits to influence the degree to which empathy-related differences between conditions translate into differences in social alignment.

## Method

### Participants

To examine the required sample size, a power analysis ^77^ was conducted for paired t-test, power of .90, d = 0.4, and α of .05, and found that the required sample size is 55. Yet, considering the potential loss in physiological and eye-tracking signals and computational model, we recruited 75 participants, among which 2 were excluded (1 due to not understanding the instructions and the other one falling asleep during the task), leading to 73 participants in total (45 females) with average age of 26.041 (SD = 4,973). Exclusion criteria was the existence of neurological or physiological disorders or current usage of medicine. All participants had normal or corrected to normal vision. The study was conducted in accordance with the Declaration of Helsinki, and was approved by the Ethics committee of Shahid Beheshti university. Written informed consents were obtained from the participants.

### Procedure

After volunteering to participate, participants received a link to an online survey administered via Porsline. The survey included demographics and health related questions, as well as the Toronto Alexithymia Scale-20 (TAS-20) and the Autism-Spectrum Quotient (AQ). Participants who met the eligibility criteria were invited to the laboratory within the following three days to take part in the experimental session. Upon arrival, participants were seated comfortably in front of a laptop with an eye-tracking system attached to it. The task instructions were presented verbally, and participants were given sufficient time to familiarize themselves with the procedure and relax before the experiment began. They were informed that, during the task, they would first view pictures of previous participants who had allegedly taken part in the study and received electrical shocks. For each picture, they were instructed either to empathize with the person shown or to remain objective. Participants were then told that, on each block, they would perform a random dot task together with one of these previous participants. Importantly, they were informed that the target participants were not present during the current experimental session, but had completed the same random-dot task in an earlier session. Thus, after providing their initial response on each trial, participants would be shown the target participant’s previous response to the same stimulus. They would then have the opportunity to revise their own response based on both their initial decision and the displayed response of the target participant. Following the instructions, electrodes were attached, and participants placed their chin on a chin rest positioned approximately 65 cm from the monitor. After eye-tracker calibration, participants began the empathy for pain and conformity task.

### Stimuli

A previously validated dataset of facial pain expressions was used in the present study. Prior work has demonstrated the validity of this dataset for eliciting empathic responses and its suitability for pain-related research ^6,78,79^. Four stimuli were selected from the dataset, consisting of two male and two female faces. Each image was overlaid with an oval mask (Figure 1).

**Figure 1.**
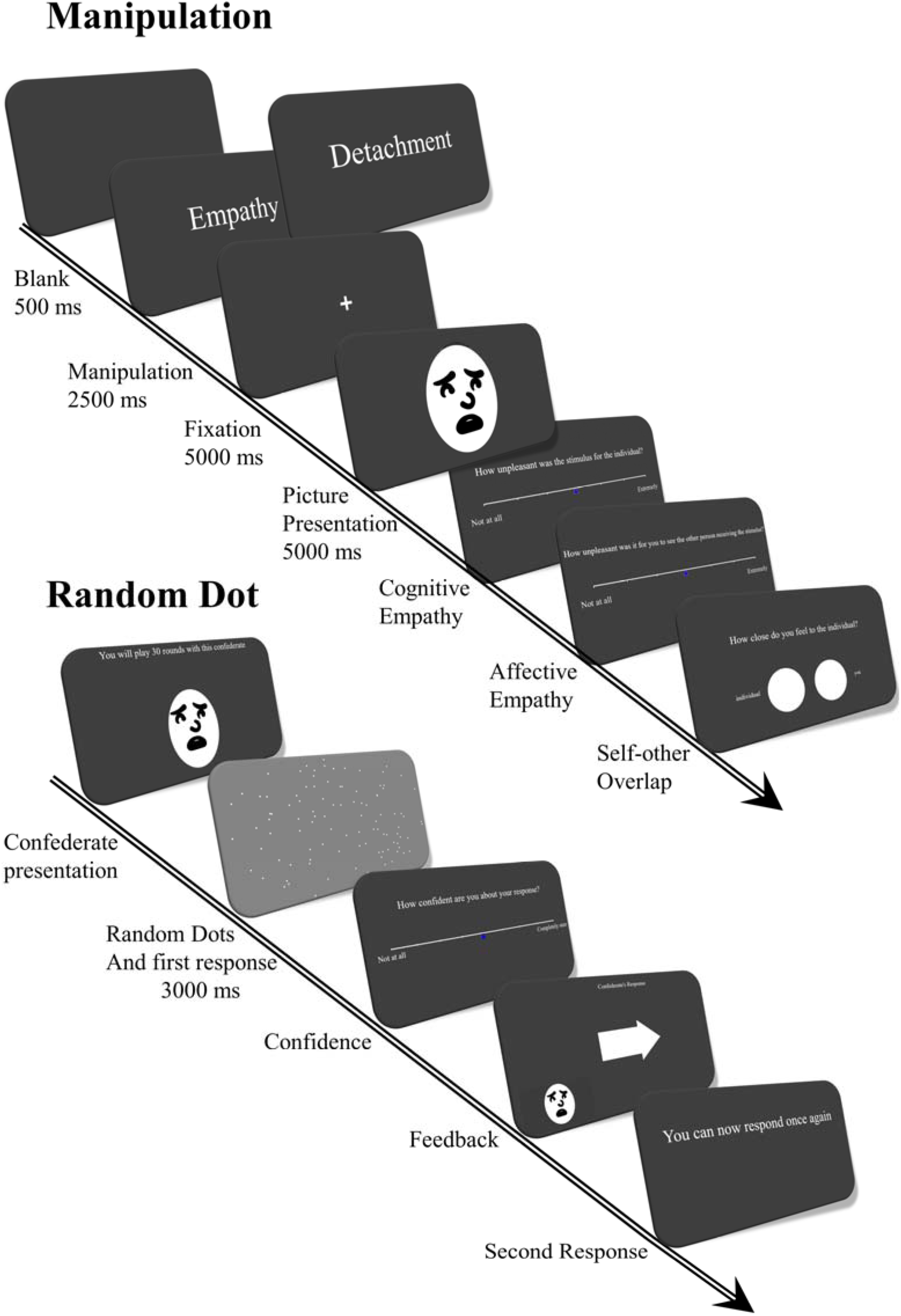
Schematic Diagram of the Empathy for Pain and Conformity Task. Note. Facial expressions were replaced by cartoon illustrations in accordance with repository restrictions.

### Empathy for Pain and Conformity Task

The task was divided into three parts of empathizing, memory check, and random dot game. During the empathizing part, participants were first presented with a 5000 milliseconds (ms) blank screen, followed by a cue presenting for 2500 ms depicting either empathy or detachment, which guides the participant regarding their behavior to the picture presented afterwards. After a 5000 ms presentation of a fixation cross, a picture depicting an individual expressing pain was shown to the participant for 10000 ms. Participants were told that the pictures had been taken from previous participants who had taken part in the study and received electrical shocks via a Digitimer DS5 isolated bipolar constant current simulator. As in previous studies ^6,7,80–83^, in case of empathy cue, participants were asked to empathize with the individual in the picture, while for the detachment, they were asked to stay objective and detached. Next, participants were presented with cognitive empathy question (How unpleasant was the stimulus for the individual?), affective empathy question (How unpleasant was it for you to see the other person receiving the stimulus?) ^7,84,85^, and Inclusion of Other in the Self scale (IOS ^86^) as a measure of interpersonal distance, during which participants were presented with two circles, that they could move toward each other or move further from each other, asking to put them in a place where represents how close they think they were to the person in the picture. None of these questions had time limit, yet considering the importance of time of physiological signals, if they were answering any question in less than 5 seconds, they were presented with a blank screen up to the difference between 5 seconds and the response time.

To ensure that participants remembered the previously seen targets’ pictures, they were then presented with a memory check, during which each picture was presented once again. Participants were asked to indicate, using the right and left buttons, whether they had empathized with the target in the picture. In case they responded wrong, they were presented with the picture once again using the same procedure as in the empathizing part, yet without responding to cognitive empathy, affective empathy and IOS questions (12.5% of trials). Next, they started the random dots phase, during which they were told that they would play 30 rounds with each of the previously seen individuals. Participants were told that these individuals have previously attended the study and their responses are recorded and would be used here. Participants were first presented with a picture of the individual they would play with, displaying a neutral facial expression. They were told that they would play 30 rounds with this individual and were asked to press the Shift key when they were ready to start the game. They had as much time as they wanted to start the game, yet the picture disappeared after 10 seconds. Within each trial an array of random dots moving either to right or left was presented to the participant. Participants were asked to indicate whether the dots were moving to the right or to the left by pressing the corresponding right or left key. Each display included 100 dots with a size of 10 pixels, moving in the direction of −15 to 15 or 165 to 195 in spherical coordinates at a speed of 10.8 pixel per frame, corresponding to 0.01 of the monitor height. Dot motion coherence was set to 15% and dots had an infinite lifetime. Participants were asked to respond as fast and accurate as possible. Dots were presented for 3 seconds and then disappeared, but responses were recorded without time limit. Next, participants were presented with a visual scale, where they had to report their confidence about their response (How confident are you about your response) using a 9-point Likert scale, ranging from not at all, to completely sure. Next, the confederate’s response was shown to the participant using an arrow on the middle of the screen pointing to the right or left, while the individual’s picture was also presented on the bottom left side of the screen. These proposals were coded so that they were aligned with participants’ choices on half of the trials and misaligned on the other half, with aligned and misaligned trials randomly intermixed throughout the task. Next, participants were given the opportunity to decide once again about their response, based on their previous decision and the response they saw from the confederate. Trials with incongruency between participants’ responses and confederate’s responses were used for the conformity ^87–89^, to examine how often participants change their opinion, when such incongruency exists. The task consisted of 4 blocks of 30 trials in total, among which 60 were related to high empathy and 60 were related to low empathy condition. For each participant, high- and low-empathy blocks were interleaved, and the starting condition was randomly assigned. The task was coded in Psychopy.

### Drift Diffusion Modeling

To model conformity-related changes in participants’ choices and reaction times, we used drift-diffusion modeling. The diffusion model was fit using fast-dm ^90,91^. We modeled participants’ second responses on trials in which their initial response was incongruent with the displayed prior response attributed to the target/confederate. Thus, the modeled responses captured participants’ decisions after viewing the target/confederate’s ostensible prior response. Given the forced-choice structure of the task, all participants provided the same number of responses. Relative starting point, reflecting bias, drift rate, the standard deviation of these parameters, alongside threshold separation were allowed to vary across empathy conditions, while the duration of non-decisional processes and the speed of response execution were considered to be equal across conditions. This model was theoretically motivated by our experimental design and research question. Yet, we additionally evaluated nine alternative model specifications and estimated the Akaike Information Criterion (AIC) for each model. The theoretically motivated model yielded the lowest AIC, indicating the best relative fit to the data (see Supplementary Materials). Furthermore, following previous studies ^92–95^, one participant was excluded since the observed data deviated significantly from the model-predicted data. Importantly, excluding this participant did not alter the pattern of results (All results including this participant are reported in the Supplementary Materials).

The estimated computational parameters capture separable cognitive mechanisms involved in conformity. Response bias is the most widely used computational index of conformity in decision-making studies ^50,51^, as it reflects a judgmental tendency to shift one’s response toward the feedback, opinions, or decisions of others. In contrast, drift rate captures the efficiency of evidence accumulation after social feedback, reflecting how individuals extract information from others’ responses and integrate it with their prior judgment when making a second decision. Finally, threshold separation reflects response caution or decisional conservativeness, indexing the amount of evidence required before committing to a choice.

### Eye-Tracking Measurement and Analysis

Eye movement data were recorded using an SMI-Red-250, with a 13-point calibration followed by a 5-point validation procedure. Eye-tracking data were analyzed for two different task epochs. First, the empathy manipulation phase, during which participants viewed the emotional facial expressions of targets, and the random dot phase, during which participants viewed the neutral facial expression of the target/confederate while waiting for the game to begin. Given the importance of eyes and mouth in empathy and facial expressions of pain, these two regions were considered as areas of interest (AOIs). As in previous studies ^53,96^, both regions were defined using rectangles extending to the sides of the face, where for eyes the rectangle was spanned from the top of the eyebrows to the equivalent distance below the eyes and for mouth it extended from the top to the bottom of the mouth. In cases where tracking ratio was below 70%, the trial was excluded (2.898% of trials in manipulation phase and 11.772% of trials in random dot phase), and in cases where both trials of a same condition had tracking ratio of below 70% the participant was excluded from analysis (4 participants in the manipulation phase, and 7 participants in the random dot phase). Fixation time and revisits were calculated for each area of interest. For epochs related to the random-dot phase, eye-tracking measures were normalized by stimulus presentation duration to account for differences in epoch length; specifically, values were divided by presentation duration in seconds. Eye-movement data were analyzed using BeGaze and MATLAB.

### Physiological signals recording and analysis

Electrocardiogram (ECG) and Skin conductance activity (SC) were recorded using bioline acquisition system at 250 Hz. ECG electrodes were placed on the wrist and clavicle, and SC electrodes were placed on index and middle finger of the non-dominant hand. ECG signal was band passed in range of 1 to 50 Hz, and detrended, and R-peaks were first defined automatically, based on the upper and lower threshold and minimum and maximum IBI, and then were checked and corrected manually. Duration of the stimulus/picture presentation within the empathy manipulation was considered as epoch, and RMSSD, and PNN-50 as the percentage of consecutive IBIs with at least 50 milliseconds difference were extracted. Skin conductance signal was first low pass filtered at 2 Hz, then tonic component was extracted using a moving minimum filter with a window size of 30s, and them smoothing using a Gaussian kernel with the same window size. The phasic component was extracted applying a first order high pass Butterworth filter of 0.5 Hz. Average Tonic and Phasic signals were measured for each condition and the difference between the conditions were calculated and used in the study. All analyses were done using PhysioData toolbox and MATLAB. Due to technical problems signals from nine participants could not be recorded, and trigger recordings were incomplete or corrupted for two additional participants. In addition, three participants were excluded from ECG analysis due to excessive noise. Also, skin conductance data were missing for two participants, resulting in final samples of 59 participants for the ECG analysis and 60 participants for the skin conductance analysis.

### Toronto Alexithymia Scale (TAS-20)

The TAS-20 ^75^ assesses alexithymia across three facets of difficulty identifying feelings, difficulty describing feelings and externally oriented thinking. Items are responded on a 5-point Likert scale. The validated Farsi version used in the present study ^97,98^ has shown good internal consistency, with reliability estimates ranging from .72 to .82. Permission to use the TAS-20 was obtained from the copyright holders.

### Autism Quotient (AQ)

Autistic traits were measured using the 28-item abridged version of the AQ ^76,99^. The AQ assesses autistic traits across five facets, as well as providing a total score. Each item is responded to on a 4-point Likert scale. The validated version used in the present study has shown acceptable reliability of .61 ^100^.

### Statistical Analyses

Paired-samples t-tests were used to assess the effectiveness of the empathy manipulation by comparing cognitive empathy, affective empathy, and self–other overlap between the high-and low-empathy conditions. Paired-samples t-tests were also used to compare the two empathy conditions on the main conformity-related model parameters, namely bias, threshold separation, and drift rate. In line with previous work using difference-score approaches ^55,101–103^, we examined whether condition-related changes in conformity were associated with condition-related changes in physiological and eye-movement measures. To this end, we computed difference scores for each variable by subtracting values in the low-empathy condition from those in the high-empathy condition, and then tested the associations between these difference scores. For conformity-related parameters, this yielded difference scores such as bias_diff_, calculated as bias in the high-empathy condition minus bias in the low-empathy condition. The same procedure was applied to all physiological and eye-movement measures. For instance, RMSSD_diff_ was calculated as RMSSD in the high-empathy condition minus RMSSD in the low-empathy condition. We then examined correlations between condition- related changes in conformity-related parameters and condition-related changes in physiological and eye-movement measures. Because physiological and eye-movement measures deviated from normality, Spearman’s rank-order correlations were used for these analyses. We also tested whether condition-related changes in conformity-related parameters were associated with individual-difference measures, namely TAS-20 and AQ scores.

Given that this is, to our knowledge, the first study to examine how empathy modulates conformity and how this effect is related to physiological responses, eye movements, and individual differences, our primary analyses focused on bivariate associations between conformity-related difference scores and each physiological, eye-movement, and individual-difference measure separately. This approach allowed us to characterize the pattern of associations between condition-related changes in these measures and condition-related changes in conformity-related decision parameters. Because these analyses were exploratory, we report uncorrected p-values and interpret them cautiously, with emphasis on effect sizes and the consistency of the observed pattern rather than on individual significance tests. To further account for potential shared variance among predictors, we also conducted exploratory multiple linear regression analyses in which conformity-related difference scores were predicted jointly by physiological measures and, in separate models, by eye-movement measures. These regression results are reported in the Supplementary Materials. In addition, because drift rate and threshold separation were exploratory variables of interest, the results concerning associations between changes in these variables and alterations in physiological and eye-tracking measures are presented in the Supplementary Materials.

## Results

### Manipulation Check

To make sure that manipulation has affected participants empathy, we compared cognitive empathy, affective empathy and social distance across conditions, using paired t-test. Results showed significantly higher cognitive, *t*(71) = 2.366, *p* = .021, and affective empathy, *t*(71) = 7.165, *p* < .001, as well as reduced social distance, *t*(71) = 5.870, *p* < .001, in high empathy condition compared to low empathy condition (Table 1 and Figure 2).

**Figure 2:**
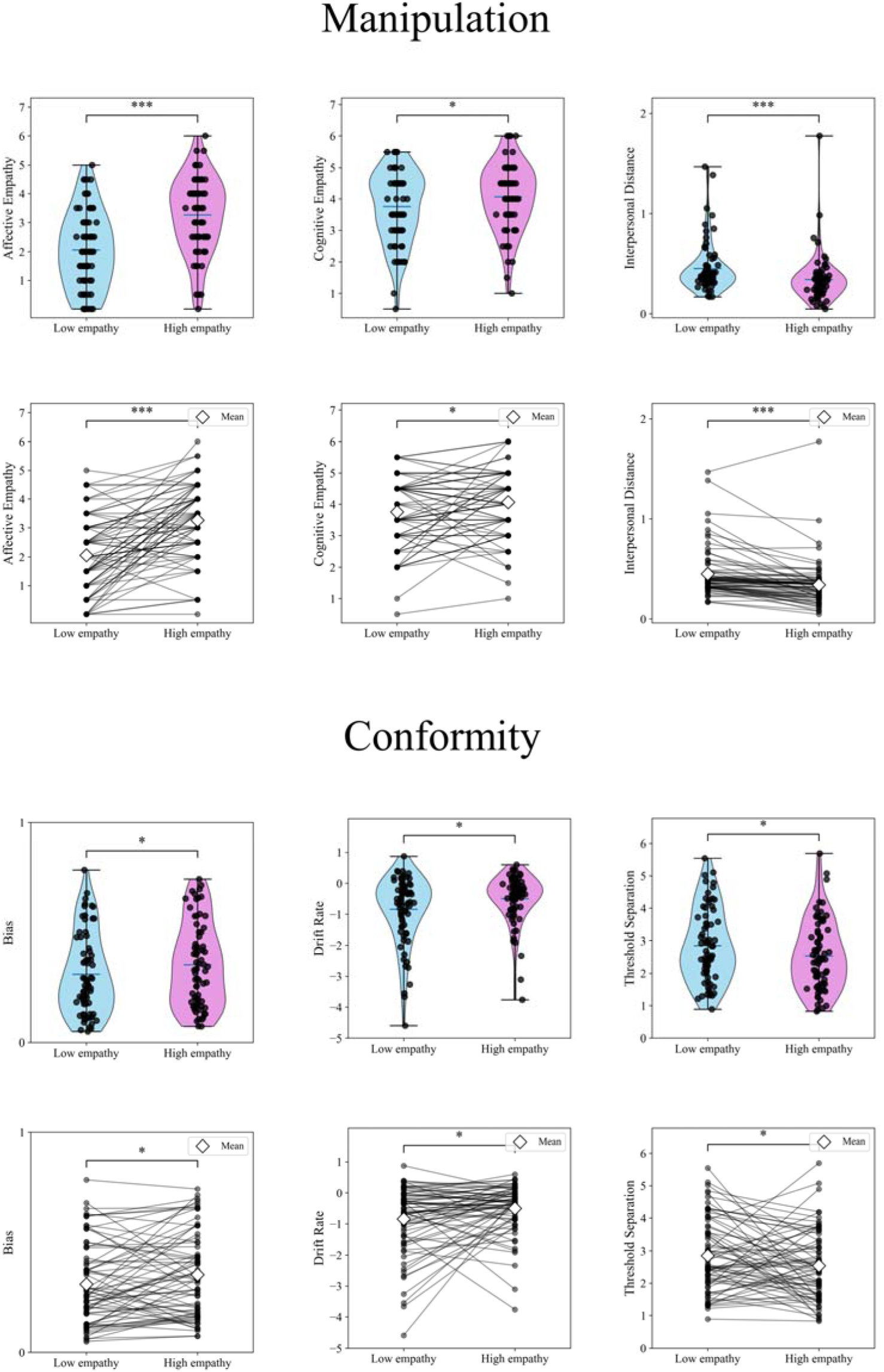
Manipulation Check and Conformity Across the Task. Notes. *p < 0.05, **p < 0.01, ***p < 0.001

**Figure 3.**
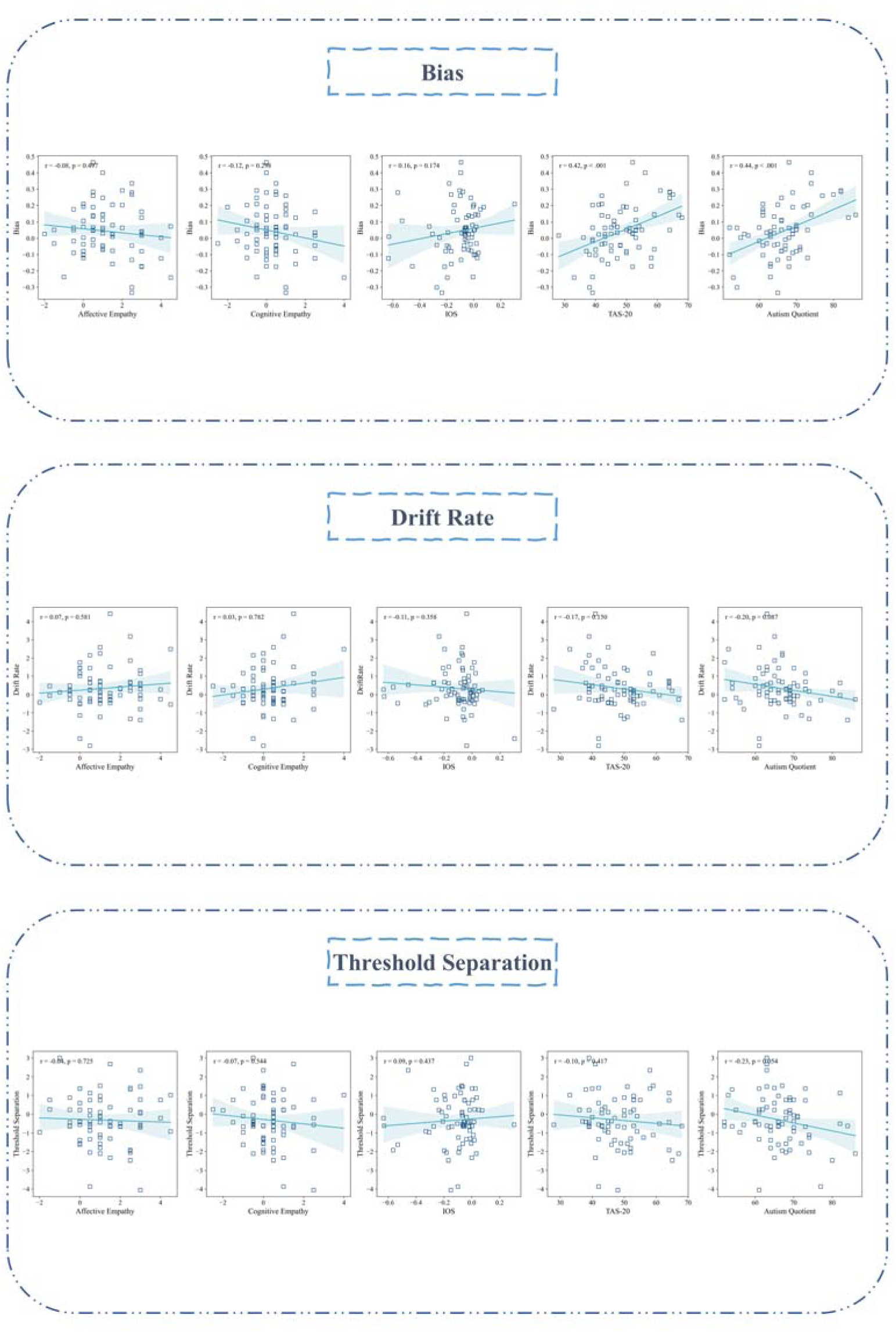
Associations between computational measures of conformity, state-empathy and self–other overlap indices, and questionnaire-based individual difference measures. Notes. *p < 0.05, **p < 0.01, ***p < 0.001

**Table 1.**
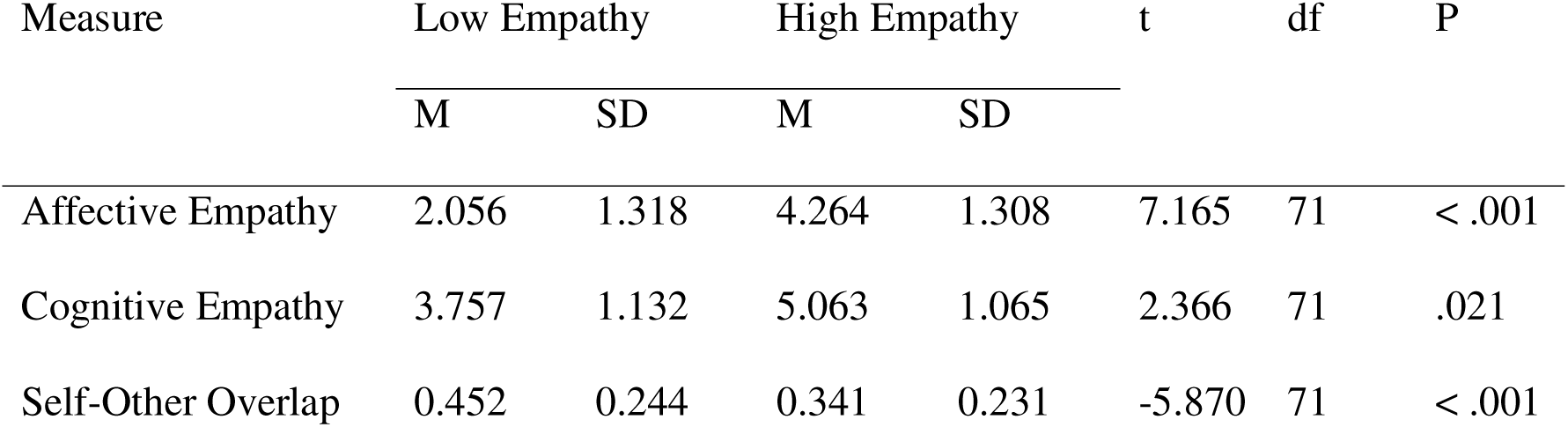
Differences in Task-Derived Cognitive Empathy, Affective Empathy, and Self–Other Overlap Across Conditions.

| Measure | Low Empathy |  | High Empathy |  | t | df | P |
| --- | --- | --- | --- | --- | --- | --- | --- |
|  | M | SD | M | SD |  |  |  |
| Affective Empathy | 2.056 | 1.318 | 4.264 | 1.308 | 7.165 | 71 | < .001 |
| Cognitive Empathy | 3.757 | 1.132 | 5.063 | 1.065 | 2.366 | 71 | .021 |
| Self-Other Overlap | 0.452 | 0.244 | 0.341 | 0.231 | -5.870 | 71 | < .001 |

### Conformity

Using relative starting point as the judgmental conformity bias measure, we found a higher level of conformity (higher bias to change the mind and make a decision in line with the confederate) in high empathy condition compared to low empathy condition, t(71) = 2.344, p = .022. Although exploratory, we also found a significantly higher drift rate, t(71) = 2.559, p = .013, and lower threshold separation, t(71) = 2.014, p = .048, in high empathy condition compared to low empathy condition. Yet, we didn’t find any relationship between judgmental bias and affective empathy, r(71) = -.081, p = .497, cognitive empathy, r(71) = -.124, p = .298, or interpersonal distance, r(71) = .162, p = .174 (Tables 2, 3; Figures 2, 3).

**Table 2.** Differences in Computational Measures of Conformity Across Conditions.

| Measure | Low Empathy |  | High Empathy |  | t | df | P |
| --- | --- | --- | --- | --- | --- | --- | --- |
|  | M | SD | M | SD |  |  |  |
| Bias | 0.309 | 0.183 | 0.352 | 0.189 | -2.344 | 71 | .022 |
| Threshold Separation | 2.849 | 1.116 | 2.533 | 1.061 | 2.014 | 71 | .048 |
| Drift Rate | -0.847 | 1.078 | -0.501 | 0.784 | -2.559 | 71 | .013 |

**Table 3.** Associations Between Computational Measures of Conformity and Task-Derived Affective Empathy, Cognitive Empathy, and Self–Other Overlap.

| Measure | Bias | Threshold Separation | Drift Rate |
| --- | --- | --- | --- |
| Affective Empathy | -.081 | -.042 | .066 |
| Cognitive Empathy | -.124 | -.073 | .033 |
| Self-Other Overlap | .162 | .093 | -.110 |

### Association Between Empathy-Related Conformity Changes and Eye-Movement Alterations

For stimuli presented during the empathy manipulation phase, a significant positive correlation was found between judgmental conformity bias and fixation duration to the mouth, r(67) = .267, p = .028. On the other hand, a significant positive correlation was found between bias and fixation duration, r(64) = .335, p = .006, as well as revisit, r(58) = .295, p = .023, in the eyes region for the stimuli presented in the random dots phase (Table 4, Figure 4).

**Figure 4.**
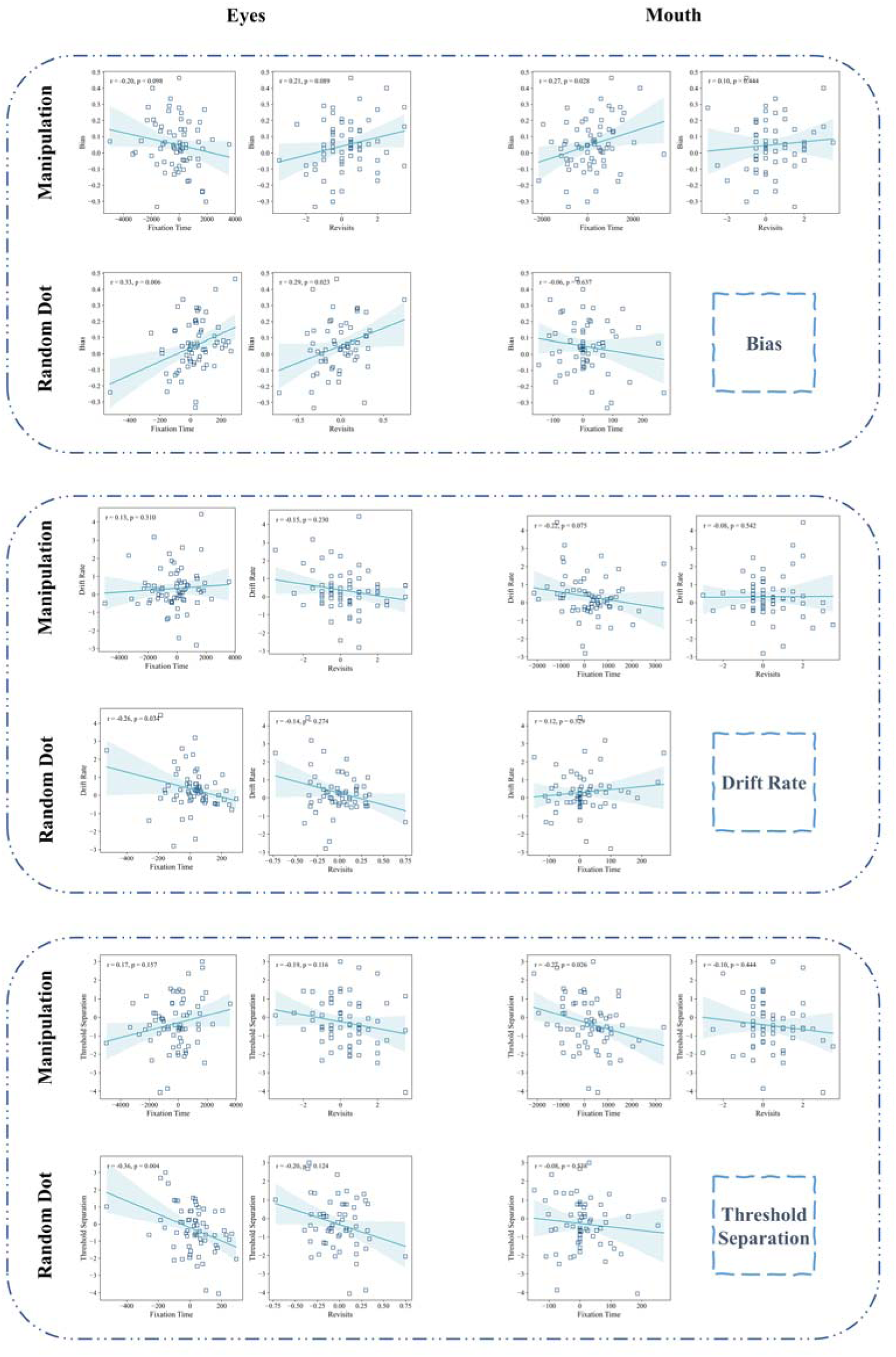
Associations between computational measures of conformity and eye-movement indices across distinct task phases. Notes. *p < 0.05, **p < 0.01, ***p < 0.001

**Table 4.** Associations Between Computational Measures of Conformity and Eye-Movement Indices Across Distinct Task Phases.

| Measure | Bias | Threshold Separation | Drift Rate |
| --- | --- | --- | --- |
| Manipulation Phase |  |  |  |
| Eye Fixation Time | -.202 | .174 | .125 |
| Eye Rivisits | .209 | -.194 | -.149 |
| Mouth Fixation Time | .267* | -.270* | -.217 |
| Mouth Revisits | .100 | -.100 | -.080 |
| Random Dot Phase |  |  |  |
| Eye Fixation Time | .335** | -.356** | -.264* |
| Eye Revisits | .295* | -.203 | -.145 |
| Mouth Fixation Time | -.060 | -.078 | .123 |

### Association Between Empathy-Related Conformity Changes and ECG and Skin Conductance Alterations

No significant correlation was found between conformity and RMSSD, r(58) = -.097, p = .463, or PNN-50, r(58) = -.142, p = .284. However, we found a significant negative correlation between conformity and average of phasic component of skin conductance, r(59) = -.319, p = .013, but not the tonic component, r(59) = -.138, p = .294 (Table 5, Figure 5).

**Figure 5.**
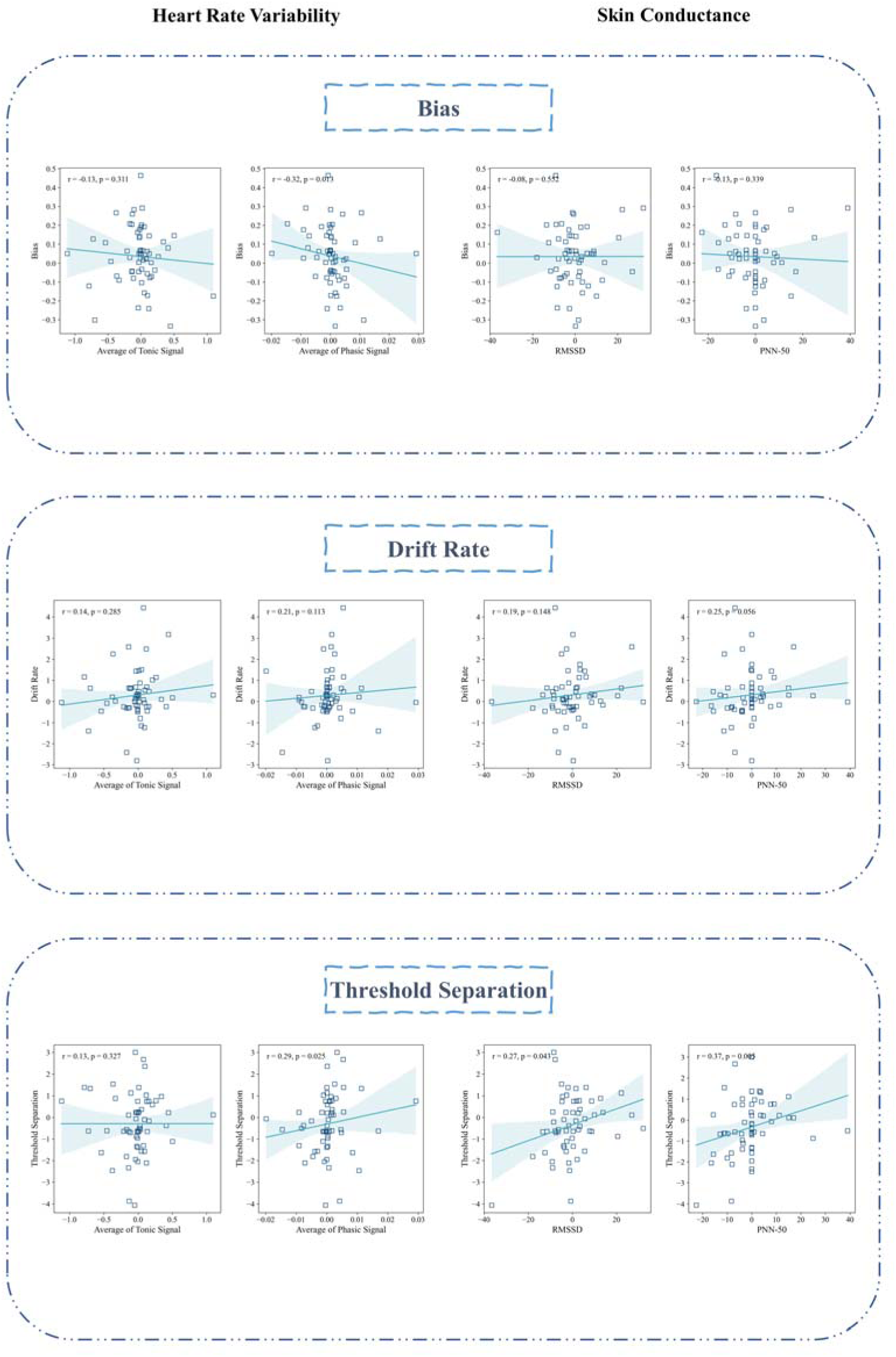
Associations Between Computational Measures of Conformity and Physiological Measures Across Distinct Task Phases. Notes. *p < 0.05, **p < 0.01, ***p < 0.001

**Table 5.**
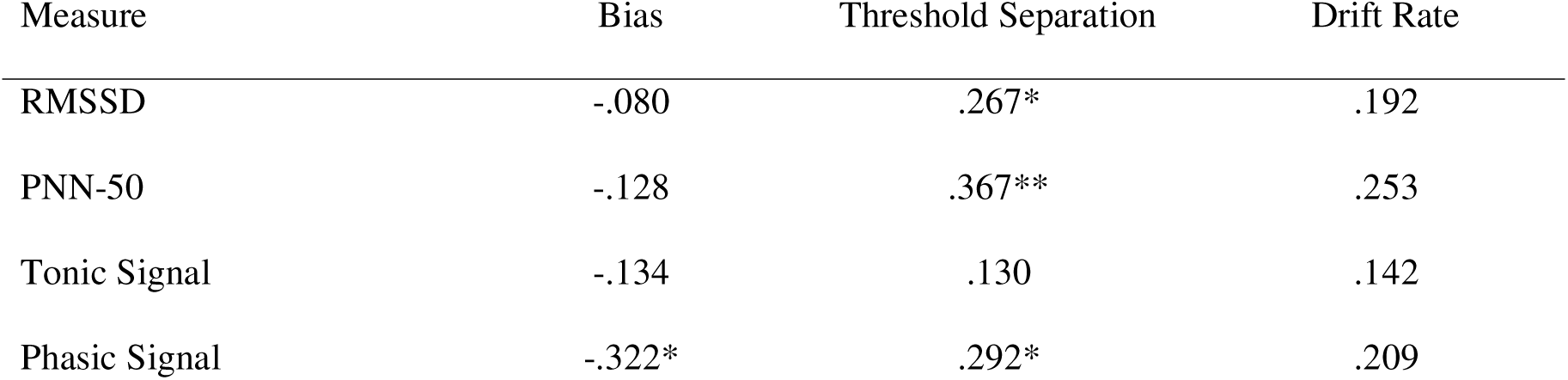
Associations Between Computational Measures of Conformity and Physiological Measures Across Distinct Task Phases.

### Alexithymia and Autistic Traits

Conformity was positively correlated with both alexithymia, r(71) = .423, p < .001, and autistic traits, r(71) = .441, p < .001 (Table 6, Figure 3).

**Table 6.**
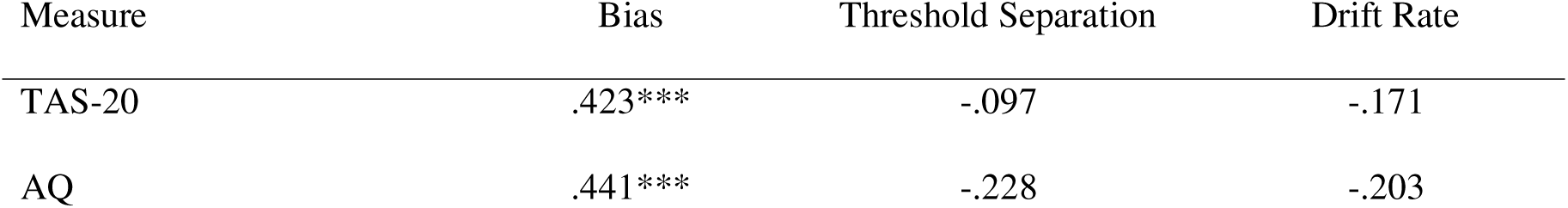
Associations Between Computational Measures of Conformity and Individual Differences in Alexithymia and Autism Traits.

### Demographics and Conformity

There were no significant sex differences in bias, *t*(70) = −1.023, *p* = .310, drift rate, *t*(70) = 1.593, *p* = .116, or threshold separation, *t*(70) = 0.397, *p* = .692. In addition, age was not significantly associated with any conformity-related parameter, including bias, *r*(70) = −.090, *p* = .450, threshold separation, *r*(70) = −.038, *p* = .751, or drift rate, *r*(70) = .094, *p* = .431.

Thus, neither sex nor age appeared to account for individual differences in conformity-related decision parameters.

## Discussion

In the present study, we investigated whether empathy promotes social alignment and conformity, and at which levels of decision-making such an effect emerges. More specifically, we examined whether empathic engagement with another person alters subsequent decisions in the direction of that person’s responses, and whether this effect is associated with visual attention to different facial regions, physiological responses during empathy, and individual differences in alexithymia and autistic traits. Our findings provide evidence that empathy does not merely increase emotional resonance and self-other overlap, but can also shape the computational mechanisms through which people make social decisions. Participants showed greater conformity toward the target of empathy, and this conformity was expressed across multiple levels of the decision process, including shifts in bias toward the empathized target, changes in the speed of evidence accumulation, and alterations in the threshold for updating one’s response based on the target’s feedback.

These findings are in line with the theory that affective alignment can facilitate social-cognitive alignment ^16^, highlighting the need to examine how alignment at one level of social processing may cascade into, shape, or reinforce alignment at other levels. Earlier studies have also shown that empathy can alter cognition in ways that reduce perceived distance from the target, including both physical and temporal distance ^6,7^. The present results suggest that this alignment extends beyond perceptual or representational closeness. Empathy appears to bias the decision-making processes toward the person with whom one empathizes, producing a form of social alignment that is expressed as conformity. In this sense, empathy may not only make another person feel closer, but may also make their judgments more influential.

One possible interpretation of this effect comes from predictive-coding accounts of social cognition ^12,14,104^. Empathy makes discrepancies between one’s own decision and the other person’s decision more salient. A mismatch between the participant’s response and the empathized target’s response may therefore generate a socially meaningful prediction error, motivating the participant to update their subsequent decisions in a direction more attuned to the other person. This interpretation is also consistent with evidence that empathy and prosocial concern can increase the priority assigned to another person’s welfare or outcomes ^105,106,106,107^, as well as with findings suggesting that learning signals can become more similar when individuals make decisions for themselves and for others ^108,109^. Thus, empathy-induced conformity may reflect not only a tendency to minimize discrepancies between self-generated and other-generated information, but also a shift in the motivational priority assigned to the empathized target. However, this interpretation should be treated cautiously, because we did not observe a significant association between IOS scores and conformity. Thus, our findings cannot be taken as direct evidence that self–other overlap mediated the effect of empathy on conformity. One possibility is that the IOS, while useful and widely used, may not be sensitive enough to capture trial-level or context-dependent changes in self–other representation. Previously it is suggested that IOS may not tap into the emotional and implicit aspects of social alignment ^110^. Future studies using more fine-grained behavioral, computational, or neural measures of self–other overlap may be better able to determine whether empathic conformity is driven by increased representational overlap, altered prediction-error processing, or greater prioritization of the empathized target.

Importantly, the pattern observed here also suggests that empathy-induced conformity differs from other forms of social influence. For example, greater conformity to ingroups relative to outgroups has been reported primarily in the speed of evidence accumulation rather than in judgmental bias ^44^. Conformity to a majority has been associated with stronger bias, but also with greater conservativeness ^50^. Similarly, norm conformity has been linked to faster evidence accumulation, whereas changes in bias appear to be less consistent and may depend on the salience of the norm ^51^. In contrast, empathy-related conformity in the present study was expressed across several components of the decision process. This suggests that empathy does not operate as a simple social cue, group signal, or majority-pressure effect. Rather, it may engage a broader set of affective, attentional, physiological, and inferential processes that jointly tune the individual toward the empathy target.

This distinction is further supported by the contrast between empathy and stress. Previous work has shown that stress can reduce conformity ^111^, whereas in the present study empathy increased conformity. Moreover, stress-related arousal, indexed by cortisol, has not consistently explained conformity effects, whereas we observed associations between empathy-related conformity and physiological indices. This suggests that the present effects cannot be reduced to general arousal or emotional intensity. Instead, empathy may represent a more specific affective-interpersonal state that changes how social information is weighted during decision-making.

A further contribution of the present study concerns the role of visual attention. Previous work has shown that conformity can itself shape attention toward conformity-relevant stimulus features ^112^. Our findings extend this idea by suggesting that visual attention before and during perceptual decision-making may influence the degree and type of conformity that subsequently emerges. Attention to the eyes and mouth has repeatedly been implicated in empathy and emotion perception, particularly in the perception of pain ^6,7,53,54^. However, our results suggest that the meaning of gaze to specific facial regions depends on both the task phase and the computational component of conformity being considered. During the empathy phase, greater attention to the mouth was associated with stronger bias toward the empathized target. This may reflect the specific emotional content of the stimuli. In painful facial expressions, the mouth may carry particularly salient information about distress, especially when the eyes are partially closed due to pain, as was the case in our stimuli. Thus, looking longer at the mouth may have strengthened affective engagement with the target, thereby increasing the tendency to align with that person’s subsequent decisions. In contrast, during the second confrontation, immediately before the random-dot task, greater attention to the eyes was associated with bias. At this stage, the task context may have shifted from affective engagement to social coordination, trust, and performance monitoring. The eyes may be especially relevant in such contexts, because they convey information about agency, intention, confidence, and social relevance ^113–117^. At the same time, the attentional findings should not be overinterpreted. In our stimuli, the pain expressions often involved tightened eyes, whereas the neutral expressions shown before the random-dot task included open eyes. Therefore, differences in gaze to the eyes and mouth may partly reflect differences in stimulus salience rather than purely psychological differences between empathy and decision preparation.

Moreover, visual attention showed different relationships with different decision components. For example, attention to the eyes in the random dot phase was related to higher bias, but slower evidence accumulation. Thus, attention to the social partner’s eyes may initially bias participants toward the partner’s response, while subsequently competing with the accumulation and integration of task-relevant sensory evidence and feedback. Another possibility is that participants who remained more socially engaged with the target adopted a more cautious or socially monitored decision strategy. Future studies should manipulate facial-region visibility or task structure more directly to disentangle these possibilities.

The physiological findings further indicate that empathy-induced conformity is not a unitary phenomenon. Bias was not reliably associated with HRV measures, whereas higher HRV reflected in both RMSSD and PNN-50 was related to greater conservativeness in the high-empathy relative to the low-empathy condition. In addition, higher RMSSD was associated with faster evidence accumulation. These findings indicate that different computational components of conformity may depend on partially distinct physiological processes. Because RMSSD and related HRV indices are often interpreted as markers of parasympathetic regulation, the association between HRV and conservativeness may reflect greater regulatory capacity during empathic engagement ^118–120^. Participants with stronger parasympathetic regulation may have required more evidence before changing their decisions in line with the empathized target, suggesting a more regulated form of social alignment rather than impulsive conformity. The positive association between RMSSD and evidence accumulation is particularly interesting. Rather than slowing decisions, stronger parasympathetic regulation may have enabled more efficient processing of task-relevant evidence under empathic conditions. In other words, physiological regulation may support a form of empathy-related conformity that is not merely emotionally driven, but cognitively organized. This interpretation is consistent with broader accounts linking vagally mediated HRV to social engagement, emotion regulation, and adaptive interpersonal responding ^120–122^. Nevertheless, because physiological measures were correlational in the present study, future work should test whether autonomic regulation plays a causal role in shaping different components of empathic conformity.

Finally, we found that individual differences in alexithymia and autistic traits moderated the relationship between empathy and conformity. Previous studies have consistently linked both alexithymia and autistic traits to differences in empathy ^7,54,56,123–126^, prediction-error processing ^66,68,69,127,128^, physiological regulation, and emotional responding ^64,65,129^. From a predictive-coding perspective, these traits may be associated with atypical weighting of prediction errors or reduced flexibility in regulating responses to social discrepancies. This may help explain why the effects of empathy on judgmental bias, as well as its association to alterations in visual attention, were more pronounced among individuals with higher alexithymia or autistic traits. The existing literature on conformity and social alignment in autism and alexithymia is mixed. Some studies suggest increased conformity ^130,131^, whereas others suggest reduced conformity ^132–135^, or no reliable differences ^136,137^. Our findings may contribute to understanding this inconsistency by showing that conformity is not a single process. Depending on whether conformity is expressed as bias, evidence accumulation, threshold adjustment, or feedback-based updating, individual-difference effects may vary.

Thus, alexithymia and autistic traits may not simply increase or decrease conformity; rather, they may alter the specific mechanisms through which empathic and social information influence decision-making.

In conclusion, the present study shows that empathy can increase social alignment and conformity, and that this effect is expressed across multiple mechanisms of decision-making. Empathy shifted participants toward the empathized target not only at the level of overt choice, but also through changes in bias, evidence accumulation, and decision updating.

These effects were further shaped by visual attention, physiological regulation, and individual differences in alexithymia and autistic traits. Together, the findings suggest that empathy is not only an effective response to another person’s state, but also a process that can reorganize how social information is weighted during decision-making. Empathy may therefore function as a bridge between affective resonance and social-cognitive alignment, making individuals more attuned to the thoughts, judgments, and actions of others.

### Limitations of the Study

Despite promising finding, some limitations should also be considered. First, our empathy manipulation focused on empathy for pain. Empathy, however, can involve responses to both negative and positive emotions, and the mechanisms observed here may not generalize to empathy for happiness, relief, pride, or other positive affective states. Future studies should examine whether empathy-induced conformity differs across emotional valence and motivational context. Second, we used a controlled manipulation involving a single target person and images depicting that person in pain. This design allowed us to isolate the effect of empathy under standardized conditions, but it does not capture the interactive and reciprocal nature of real-life empathy. In particular, it did not allow us to examine physiological synchrony between participant and target, which may be a central mechanism of interpersonal alignment. Dyadic designs would therefore be valuable for testing whether synchrony in physiology, gaze, or neural responses predicts subsequent conformity. Third, we relied on the IOS as a measure of self–other overlap. Although widely used, the IOS may be too coarse to detect subtle or dynamic changes in self–other representation during empathic decision-making. More sensitive measures are needed to test whether self–other overlap mediates the relationship between empathy and conformity.

## Acknowledgements

We would like to thank Mina Hosseinnezhad and Zahra Karami for their support during the participant recruitment process. We are also grateful to all participants for their time and contribution to this study.

## Resource Availability Lead author

Further information and requests for resources and reagents should be directed to and will be fulfilled by the lead contact, Khatereh Borhani.

## Materials availability

This study did not generate new unique materials.

## Data and Code Availability

All data supporting the findings of this study are deposited at the Open Science Framework repository.

All code used to analyze the data has been deposited in the Open Science Framework repository.

Any additional information required to reanalyze the data reported in this article is available from the lead contact upon request.

## Authors Contribution

SG: Conceptualization; Data Collection; Formal analysis; Investigation; Methodology; Resources; Software; Visualization; Writing—Original draft; Writing—Review & editing. MW: Writing—Review & editing. KB: Conceptualization; Formal analysis; Methodology; Project administration; Resources; Supervision; Writing—Original draft; Writing—Review & editing.

## Declaration of generative AI and AI-assisted technologies in the writing process

During the preparation of this work SG used ChatGPT solely to improve language and readability. After using these tools, the author reviewed and edited the content as needed and took full responsibility for the content of the published article.

## Supplemental information

Document S1. Tables S1–S25

