## Supplemental File for "Shared Pain, Shared Decisions: How Empathy Shapes Social Conformity Through Physiology and Visual Attention"

**Model Comparison**

In order to evaluate our model against other possible models, we fitted the data using 9 different models. Considering the fact that bias, threshold separation, and drift rate are variables of interests, they were kept varying across conditions in all models. Models differed based on whether variables of non-decisional process duration (t0), response execution duration (d), intertrial variability of non-decisional process duration (st0), intertrial variability of bias (sz), and intertrial variability of drift rate (sv) were allowed to vary across conditions, kept the same across conditions or kept as zero. All models were fitted with fast-dm using maximum likelihood and AIC was calculated for each model. Across different models, the theoretically-driven model used in the main text, showed the lowest AIC. Results are presented in Table S1.

**Speed of Evidence Accumulation and Conservativeness**

As an exploratory analysis, the relationship between changes in speed of evidence accumulation (drift rate) and conservativeness (threshold separation) with other measures were also examined. During the manipulation phase, threshold separation was negatively correlated with the fixation on the mouth, r(67) = -.270, p = .026, but not with the revisit on the mouth or on the eyes (p > .05). Also in the random dot phase, there was a significant negative correlation between fixation time on eyes region and both threshold separation, r(64) = -.356, p = .004, and drift diffusion, r(65) = -.264, p = .034. Regarding heart rate variability, threshold separation was significantly positively correlated with RMSSD, r(58) = .291, p = .026, and PNN-50, r(58) = .388, p = .002. Drift diffusion on the other hand showed no significant correlation with either RMSSD, r(58) = .203, p = .123, but a significant correlation with PNN-50, r(58) = .263, p = .044. In addition, Threshold separation showed no significant correlation with average of tonic component, r(59) = .141, p = .282, but a significant positive correlation with average of phasic component, r(59) = .285, p = .027. On the other hand, changes in drift diffusion showed no significant correlation with either changes in tonic, r(59) = .145, p = .269, or phasic component, r(59) = .206, p = .114, of the skin conductance.

**Analyses Without Exclusion**

For completeness we also report all results without excluding one participant due to poor fit of drift diffusion model. Results completely mimicked the ones with exclusion. Results are presented in Tables S2-S7.

**Mixed Effect Models**

We used mixed-effects models to investigate the effects of visual attention and physiological responses on empathy-induced conformity. Because eye-tracking measures were obtained during two distinct phases, the empathy manipulation phase and the random-dot motion phase, we analyzed these phases separately. In each model, bias was predicted by eye-tracking-derived measures of visual attention.

Consistent with the correlational analyses, fixation duration on the mouth region and revisits to the eye region during the empathy manipulation phase significantly positively predicted bias. During the random-dot motion phase, fixation duration on the eye region was the only significant positive predictor of bias. For drift rate, reflecting the speed of evidence accumulation, fixation duration on the mouth region and revisits to the eye region during the empathy manipulation phase were significant negative predictors. No eye-tracking measure from the random-dot motion phase significantly predicted drift rate. Finally, although fixation duration on the mouth region was negatively correlated with response conservativeness, indexed by threshold separation, this effect was only marginally significant in the mixed-effects model. Revisits to the eye region, however, significantly negatively predicted conservativeness. Consistent with the correlational findings, fixation duration on the eye region during the random-dot motion phase also negatively predicted conservativeness (Tables S8-S13, S17-S22). In addition, when bias using physiological signals, similar to the correlational analysis, average phasic component of skin conductance marginally significantly predicted bias. However, no significant predictor was found in models predicting threshold separation and drift rate (Tables S14-S16, S23-S25). Results remained similar in with and without excluding the participant with poor fit.

Table S1. AIC for different models fitted to the data

| Model | Parameters | k | AIC |
| --- | --- | --- | --- |
| Model 1 | t0: inv, st0: inv, d: inv, sz: dep, sv: dep | 13 | 152 |
| Model 2 | t0: dep, st0: dep, d: dep, sz: dep, sv: dep | 16 | 158 |
| Model 3 | t0: dep, st0: 0, d: dep, sz: 0, sv: 0 | 10 | 168 |
| Model 4 | t0: inv, st0: 0, d: inv, sz: 0, sv: 0 | 8 | 161 |
| Model 5: | t0: dep, st0: dep, d: 0, sz: 0, sv: 0 | 10 | 164 |
| Model 6: | t0: dep, st0: dep, d: dep, sz: 0, sv: 0 | 12 | 164 |
| Model 7: | t0: dep, st0: dep, d: inv, sz: 0, sv: 0 | 11 | 163 |
| Model 8: | t0: dep, st0: dep, d: inv, sz: dep, sv: dep | 15 | 156 |
| Model 9: | t0: dep, st0: inv, d: inv, sz: dep, sv: dep | 14 | 153 |
| Model 10 | t0: 0, st0: 0, d: 0, sz: 0, sv: 0 | 6 | 170 |

Table S2: Differences in Task-Derived Cognitive Empathy, Affective Empathy, and Self–Other Overlap Across Conditions in the Full Unexcluded Sample

| Measure | Low Empathy | | High Empathy | | t | df | P |
| --- | --- | --- | --- | --- | --- | --- | --- |
|  | M | SD | M | SD |  |  |  |
| Affective Empathy | 3,075 | 1,319 | 0,451 | 1,299 | 7.131 | 72 | < .001 |
| Cognitive Empathy | 4,747 | 1,128 | 5,062 | 1,057 | 2.467 | 72 | .016 |
| Self-Other Overlap | 0,451 | 0,243 | 0,341 | 0,230 | -5.849 | 72 | < .001 |

Table S3: Differences in Computational Measures of Conformity Across Conditions in the Full Unexcluded Sample

| Measure | Low Empathy | | High Empathy | | t | df | P |
| --- | --- | --- | --- | --- | --- | --- | --- |
|  | M | SD | M | SD |  |  |  |
| Bias | 0.307 | 0.183 | 0.351 | 0.188 | -2.404 | 72 | .019 |
| Threshold Separation | 2.884 | 1.148 | 2.554 | 1.069 | 2.125 | 72 | .037 |
| Drift Rate | -0.834 | 1.077 | -0.494 | 0.781 | -2.547 | 72 | .013 |

Table S4: Associations Between Computational Measures of Conformity and Task-Derived Affective Empathy, Cognitive Empathy, and Self–Other Overlap in the Full Unexcluded Sample

| Measure | Bias | Threshold Separation | Drift Rate |
| --- | --- | --- | --- |
| Affective Empathy | -.084 | -.029 | .073 |
| Cognitive Empathy | -.116 | -.087 | .024 |
| Self-Other Overlap | .168 | .076 | -.115 |

Table S5: Associations Between Computational Measures of Conformity and Eye-Movement Indices Across Distinct Task Phases in the Full Unexcluded Sample

| Measure | Bias | Threshold Separation | Drift Rate |
| --- | --- | --- | --- |
| Manipulation Phase | | | |
| Eye Fixation Time | -.190 | .154 | .117 |
| Eye Rivisits | .201 | -.181 | -.143 |
| Mouth Fixation Time | .260* | -.253* | -.214 |
| Mouth Revisits | .094 | -.094 | -.076 |
| Random Dot Phase | | | |
| Eye Fixation Time | .334** | -.355** | -.265* |
| Eye Revisits | .294* | -.210 | -.144 |
| Mouth Fixation Time | -.067 | -.058 | .130 |

Table S6 Associations Between Computational Measures of Conformity and Physiological Measures Across Distinct Task Phases in the Full Unexcluded Sample

| Measure | Bias | Threshold Separation | Drift Rate |
| --- | --- | --- | --- |
| RMSSD | -.097 | .291* | .203 |
| PNN-50 | -.142 | .388** | .263* |
| Tonic Signal | -.138 | .141 | .145 |
| Phasic Signal | -.319* | .285* | .206 |

Table S7. Associations Between Computational Measures of Conformity and Individual Differences in Alexithymia and Autism Traits in the Full Unexcluded Sample

| Measure | Bias | Threshold Separation | Drift Rate |
| --- | --- | --- | --- |
| TAS-20 | .422*** | -.094 | -.170 |
| AQ | .442*** | -.230 | -.204 |

Table S8. Predicting Empathy-Related Changes in Bias from Changes in Eye-Movement Indices During the Manipulation Phase

| Variable | β | SE | CI | t | df | p |
| --- | --- | --- | --- | --- | --- | --- |
| Eye Fixation Time | 0.000 | 0.000 | 0.000, 0.000 | -0.429 | 55 | 0.670 |
| Eye Rivisits | 0.036 | 0.016 | 0.004, 0.069 | 2.246 | 55 | 0.029 |
| Mouth Fixation Time | 0.000 | 0.000 | 0.000, 0.000 | 2.559 | 55 | 0.013 |
| Mouth Revisits | -0.021 | 0.018 | -0.057, 0.016 | -1.112 | 55 | 0.271 |

Table S9. Predicting Empathy-Related Changes in Bias from Changes in Eye-Movement Indices During the Random Dot Phase

| Variable | Β | SE | CI | t | df | p |
| --- | --- | --- | --- | --- | --- | --- |
| Eye Fixation Time | 0.000 | 0.000 | 0.000, 0.001 | 2.016 | 55 | 0.049 |
| Eye Revisits | 0.096 | 0.096 | -0.096, 0.289 | 1.001 | 55 | 0.321 |
| Mouth Fixation Time | 0.000 | 0.000 | -0.001, 0.000 | -1.057 | 55 | 0.295 |

Table S10. Predicting Empathy-Related Changes in Drift Rate from Changes in Eye-Movement Indices During the Manipulation Phase

| Variable | β | SE | CI | t | df | p |
| --- | --- | --- | --- | --- | --- | --- |
| Eye Fixation Time | 0.000 | 0.000 | 0.000, 0.000 | -0.015 | 55 | 0.988 |
| Eye Rivisits | -0.273 | 0.125 | -0.523, -0.023 | -2.188 | 55 | 0.033 |
| Mouth Fixation Time | 0.000 | 0.000 | -0.001, 0.000 | -1.971 | 55 | 0.054 |
| Mouth Revisits | 0.204 | 0.142 | -0.082, 0.489 | 1.429 | 55 | 0.159 |

Table S11. Predicting Empathy-Related Changes in Drift Rate from Changes in Eye-Movement Indices During the Random Dot Phase

| Variable | β | SE | CI | t | df | p |
| --- | --- | --- | --- | --- | --- | --- |
| Eye Fixation Time | -0.002 | 0.001 | -0.004, 0.001 | -1.270 | 55 | 0.209 |
| Eye Revisits | -0.718 | 0.727 | -2.176, 0.739 | -0.988 | 55 | 0.328 |
| Mouth Fixation Time | 0.002 | 0.002 | -0.002, 0.006 | 0.990 | 55 | 0.327 |

Table S12. Predicting Empathy-Related Changes in Threshold Separation from Changes in Eye-Movement Indices During the Manipulation Phase

| Variable | β | SE | CI | t | df | p |
| --- | --- | --- | --- | --- | --- | --- |
| Eye Fixation Time | 0.000 | 0.000 | 0.000, 0.000 | 1.681 | 55 | 0.098 |
| Eye Rivisits | -0.301 | 0.129 | -0.560, -0.042 | -2.327 | 55 | 0.024 |
| Mouth Fixation Time | 0.000 | 0.000 | -0.001, 0.000 | -2.064 | 55 | 0.044 |
| Mouth Revisits | 0.119 | 0.147 | -0.176, 0.414 | 0.807 | 55 | 0.423 |

Table S13. Predicting Empathy-Related Changes in Threshold Separation from Changes in Eye-Movement Indices During the Random Dot Phase

| Variable | β | SE | CI | t | df | p |
| --- | --- | --- | --- | --- | --- | --- |
| Eye Fixation Time | -0.004 | 0.001 | -0.007, -0.001 | -2.536 | 55 | 0.014 |
| Eye Revisits | -0.643 | 0.829 | -2.304, 1.019 | -0.775 | 55 | 0.442 |
| Mouth Fixation Time | -0.002 | 0.002 | -0.007, 0.002 | -1.045 | 55 | 0.301 |

Table S14. Predicting Empathy-Related Changes in Bias from Changes in Physiological Indices

| Variable | β | SE | CI | t | df | p |
| --- | --- | --- | --- | --- | --- | --- |
| RMSSD | 0.002 | 0.004 | -0.006, 0.009 | 0.465 | 51 | 0.644 |
| PNN-50 | -0.002 | 0.004 | -0.010, 0.006 | -0.600 | 51 | 0.551 |
| Average Tonic Signal | -0.095 | 0.075 | -0.244, 0.055 | -1.269 | 51 | 0.210 |
| Average Phasic Signal | -6.584 | 3.511 | -13.633, 0.464 | -1.875 | 51 | 0.066 |

Table S15. Predicting Empathy-Related Changes in Drift Rate from Changes in Physiological Indices

| Variable | β | SE | CI | t | df | p |
| --- | --- | --- | --- | --- | --- | --- |
| RMSSD | 0.008 | 0.029 | -0.050, 0.066 | 0.275 | 51 | 0.784 |
| PNN-50 | 0.007 | 0.031 | -0.056, 0.069 | 0.210 | 51 | 0.834 |
| Average Tonic Signal | 0.783 | 0.586 | -0.394, 1.961 | 1.336 | 51 | 0.187 |
| Average Phasic Signal | 33.264 | 27.594 | -22.134, 88.662 | 1.205 | 51 | 0.234 |

Table S16. Predicting Empathy-Related Changes in Threshold Separation from Changes in Physiological Indices

| Variable | β | SE | CI | t | df | p |
| --- | --- | --- | --- | --- | --- | --- |
| RMSSD | 0.016 | 0.032 | -0.049 | 0.495 | 51 | 0.623 |
| PNN-50 | 0.028 | 0.035 | -0.041 | 0.807 | 51 | 0.424 |
| Average Tonic Signal | 0.229 | 0.651 | -1.078 | 0.352 | 51 | 0.726 |
| Average Phasic Signal | 44.740 | 30.647 | -16.786 | 1.460 | 51 | 0.150 |

Table S17. Predicting Empathy-Related Changes in Bias from Changes in Eye-Movement Indices During the Manipulation Phase in the Full Unexcluded Sample

| Variable | β | SE | CI | t | df | p |
| --- | --- | --- | --- | --- | --- | --- |
| Eye Fixation Time | -0.000 | 0.000 | -0.000, 0.000 | -0.386 | 56 | 0.701 |
| Eye Rivisits | 0.035 | 0.016 | 0.003, 0.068 | 2.215 | 56 | 0.031 |
| Mouth Fixation Time | .000 | 0.000 | 0.000, 0.000 | 2.537 | 56 | 0.014 |
| Mouth Revisits | -0.020 | 0.018 | -0.057, 0.017 | -1.099 | 56 | 0.276 |

Table S18. Predicting Empathy-Related Changes in Bias from Changes in Eye-Movement Indices During the Random Dot Phase in the Full Unexcluded Sample

| Variable | β | SE | CI | t | df | p |
| --- | --- | --- | --- | --- | --- | --- |
| Eye Fixation Time | 0.000 | 0.000 | 0.000, 0.001 | 2.034 | 56 | 0.047 |
| Eye Revisits | 0.096 | 0.095 | -0.095, 0.287 | 1.010 | 56 | 0.317 |
| Mouth Fixation Time | 0.000 | 0.000 | -0.001, 0.000 | -1.070 | 56 | 0.289 |

Table S19. Predicting Empathy-Related Changes in Drift Rate from Changes in Eye-Movement Indices During the Manipulation Phase in the Full Unexcluded Sample

| Variable | β | SE | CI | t | df | p |
| --- | --- | --- | --- | --- | --- | --- |
| Eye Fixation Time | 0.000 | 0.000 | 0.000, 0.000 | -0.061 | 56 | 0.951 |
| Eye Rivisits | -0.267 | 0.124 | -0.515, -0.019 | -2.156 | 56 | 0.035 |
| Mouth Fixation Time | 0.000 | 0.000 | -0.001, 0.000 | -1.946 | 56 | 0.057 |
| Mouth Revisits | 0.201 | 0.142 | -0.083, 0.484 | 1.418 | 56 | 0.162 |

Table S20. Predicting Empathy-Related Changes in Drift Rate from Changes in Eye-Movement Indices During the Random Dot Phase in the Full Unexcluded Sample

| Variable | β | SE | CI | t | df | p |
| --- | --- | --- | --- | --- | --- | --- |
| Eye Fixation Time | -0.002 | 0.001 | -0.004, 0.001 | -1.281 | 56 | 0.206 |
| Eye Revisits | -0.722 | 0.721 | -2.165, 0.722 | -1.002 | 56 | 0.321 |
| Mouth Fixation Time | 0.002 | 0.002 | -0.002, 0.006 | 1.013 | 56 | 0.315 |

Table S21. Predicting Empathy-Related Changes in Threshold Separation from Changes in Eye-Movement Indices During the Manipulation Phase in the Full Unexcluded Sample

| Variable | β | SE | CI | t | df | p |
| --- | --- | --- | --- | --- | --- | --- |
| Eye Fixation Time | 0.000 | 0.000 | 0.000, 0.000 | 1.582 | 56 | 0.119 |
| Eye Rivisits | -0.288 | 0.130 | -0.547, -0.028 | -2.219 | 56 | 0.031 |
| Mouth Fixation Time | 0.000 | 0.000 | -0.001, 0.000 | -1.976 | 56 | 0.053 |
| Mouth Revisits | 0.113 | 0.148 | -0.184, 0.410 | 0.762 | 56 | 0.449 |

Table S22. Predicting Empathy-Related Changes in Threshold Separation from Changes in Eye-Movement Indices During the Random Dot Phase in the Full Unexcluded Sample

| Variable | β | SE | CI | t | df | p |
| --- | --- | --- | --- | --- | --- | --- |
| Eye Fixation Time | -0.004 | 0.001 | -0.007, -0.001 | -2.541 | 56 | 0.014 |
| Eye Revisits | -0.662 | 0.826 | -2.316, 0.992 | -0.802 | 56 | 0.426 |
| Mouth Fixation Time | -0.002 | 0.002 | -0.007, 0.002 | -0.991 | 56 | 0.326 |

Table S23. Predicting Empathy-Related Changes in Bias from Changes in Physiological Indices

| Variable | β | SE | CI | t | df | p |
| --- | --- | --- | --- | --- | --- | --- |
| RMSSD | 0.002 | 0.004 | -0.006, 0.009 | 0.462 | 52 | 0.646 |
| PNN-50 | -0.002 | 0.004 | -0.010, 0.005 | -0.621 | 52 | 0.538 |
| Average Tonic Signal | -0.095 | 0.074 | -0.243, 0.054 | -1.279 | 52 | 0.207 |
| Average Phasic Signal | -6.598 | 3.477 | -13.574, 0.379 | -1.898 | 52 | 0.063 |

Table S24. Predicting Empathy-Related Changes in Drift Rate from Changes in Physiological Indices

| Variable | β | SE | CI | t | df | p |
| --- | --- | --- | --- | --- | --- | --- |
| RMSSD | 0.008 | 0.029 | -0.050, 0.066 | 0.282 | 52 | 0.779 |
| PNN-50 | 0.007 | 0.031 | -0.055, 0.068 | 0.218 | 52 | 0.828 |
| Average Tonic Signal | 0.783 | 0.581 | -0.382, 1.948 | 1.348 | 52 | 0.183 |
| Average Phasic Signal | 33.303 | 27.320 | -21.519, 88.125 | 1.219 | 52 | 0.228 |

Table S25. Predicting Empathy-Related Changes in Threshold Separation from Changes in Physiological Indices

| Variable | β | SE | CI | t | df | p |
| --- | --- | --- | --- | --- | --- | --- |
| RMSSD | 0.016 | 0.032 | -0.048, 0.080 | 0.505 | 52 | 0.616 |
| PNN-50 | 0.028 | 0.034 | -0.040, 0.096 | 0.824 | 52 | 0.414 |
| Average Tonic Signal | 0.228 | 0.645 | -1.066, 1.522 | 0.354 | 52 | 0.725 |
| Average Phasic Signal | 44.802 | 30.343 | -16.086, 105.691 | 1.477 | 52 | 0.146 |
